# The Douglas Bell Canada Brain Bank Post-mortem Brain Imaging Protocol

**DOI:** 10.1101/2024.02.27.582303

**Authors:** Mahsa Dadar, Liana Sanches, Jeremie Fouqouet, Roqaie Moqadam, Zaki Alasmar, Dominique Miraut, Josefina Maranzano, Naguib Mechawar, M. Mallar Chakravarty, Yashar Zeighami

**Author notes:** **Corresponding Authors Information:** Mahsa Dadar, Yashar Zeighami, Cerebral Imaging Centre, 6875 Boulevard LaSalle, Montréal, QC, H4H 1R3.

## Abstract

Magnetic resonance imaging (MRI) is a valuable non-invasive tool that has been widely used for in vivo investigations of brain morphometry and microstructural characteristics. Postmortem MRIs can provide complementary anatomical and microstructural information to in vivo imaging and ex vivo neuropathological assessments without compromising the sample for future investigations. We have developed a postmortem MRI protocol for the brain specimens of the Douglas-Bell Canada Brain Bank (DBCBB), the largest brain bank in Canada housing over 3000 neurotypical and diseased brain specimens, that allows for acquisition of high-resolution 3T and 7T MRIs. Our protocol can be used to scan DBCBB specimens with minimal tissue manipulation, allowing for feasibly scanning large numbers of postmortem specimens while retaining the quality of the tissue for downstream histology and immunohistochemistry assessments. We demonstrate the robustness of this protocol in spite of the dependency of image quality on fixation by acquiring data on the first day of extraction and fixation, to over twenty years post fixation. The acquired images can be used to perform volumetric segmentations, cortical thickness measurements, and quantitative analyses which can be potentially used to link MRI-derived and ex vivo histological measures, assaying both the normative organization of the brain and ex vivo measures of pathology.

## Introduction

Magnetic resonance imaging (MRI) allows for non-invasive and longitudinal characterization of brain anatomy, presence of abnormalities, and neurodegeneration (Albert et al., n.d.; Frisoni et al., 2010). Structural MRIs enable volumetric quantification of brain structures as well as tumors and lesions, while diffusion-weighted and quantitative MRIs provide valuable information on microstructure and biophysical properties of the tissue (Abd-Ellah et al., 2019, p.; Concha, 2014). While MRI measures are sensitive to neurodegenerative processes and brain abnormalities, they are not specific. There is therefore a key need to supplement, validate, and characterize morphological measures and biophysical models with neurobiological information. Complementing in vivo MRI assessments, postmortem MRI can be performed at longer sessions (i.e. overnight) with higher resolutions and be linked to microscopy assessments, allowing for multi-scale investigations of the brain (Pfefferbaum et al., 2004; Schumann et al., 2001; Tendler et al., 2022). Finally, ex vivo MRI measurements can be linked to postmortem neuropathology assessments to develop potential MRI-based neuropathology biomarkers (Dawe et al., 2014; Pallebage-Gamarallage et al., 2018). For example, ex vivo transverse relaxation rate (R2*) is sensitive to the presence of amyloid plaques in aging and Alzheimer’s disease specimens (Antharam et al., 2012; Bulk et al., 2018; Meadowcroft et al., 2015). Quantitative susceptibility mapping (QSM), on the other hand, is highly sensitive and specific to the presence of local diamagnetic (e.g. myelin) and paramagnetic (e.g. iron deposits) signal sources (Langkammer et al., 2012). Combined together, multi-modal ex vivo quantitative MRI can provide a multidimensional view of the underlying tissue properties, which can advance our understanding of its pathological changes.

Neuropathology confirmation of in vivo brain MRI findings requires post-mortem assessment of the same regions through histology, which can be achieved by mapping the in vivo MRIs to the histology section images (Choe et al., 2011; Humphreys et al., 2021; Maranzano et al., 2020; Vedam-Mai, 2022; Waldvogel et al., 2006). Histology is usually performed on smaller tissue blocks that need to be mapped back to the whole brain MRI (Pallebage-Gamarallage et al., 2018). To facilitate this process, the histology blocks can be scanned (preferably using ultra-high field 7T scanners to achieve higher image resolution), providing a reliable intermediate image that can be accurately mapped to both in vivo MRIs and histology images (Goubran et al., 2015b, 2015a; Humphreys et al., 2021; Roseborough et al., 2020). Ex vivo MRIs of intact brains can also be used to accurately localize areas of interest for further histology and immunohistochemistry (IHC) assessments (i.e. presence of cerebrovascular lesions such as white matter hyperintensities, tumors, etc.) without cutting into the tissue.

In this work, we present a post-mortem MRI protocol that has been developed and optimized for the Douglas Bell Canada Brain Bank (DBCBB) specimens, allowing for data acquisition from a large number of postmortem brains without jeopardizing tissue quality for follow-up histology and immunohistochemistry assessments. We demonstrate feasibility of performing high quality post-mortem MRIs with a standard one hour protocol using a 3T human MRI scanner, as well as two higher resolution overnight protocols using the same 3T scanner as well as a 7T preclinical scanner for achieving greater levels of anatomical detail.

## Methods

### Postmortem Specimens

The DBCBB is one of the largest brain banks worldwide, housing over 3000 brains with different neurodegenerative disorders including Alzheimer’s disease, Parkinson’s disease, frontotemporal dementia, Amyotrophic lateral sclerosis (ALS), as well as diverse psychiatric conditions including schizophrenia, major depression, bipolar disorder, substance use disorders (https://douglasbrainbank.ca/).

Linked to a large relational database of demographics, family history, and neuropathology information, the DBCBB specimens provide an invaluable resource for studying the neurobiology of brain diseases. Brain specimens are collected postmortem following consent of the next of kin, according to tissue banking practices regulated by the Quebec Health Research Fund, and the Guidelines on Human Biobanks and Genetic Research Databases (Pidsley et al., 2014).

After reception at the DBCBB, hemispheres are separated by a sagittal cut in the middle of the brain, brainstem, and cerebellum. One unsliced hemisphere (right or left, in alternation) is fixed in 10% neutral buffered formalin (in 7-liter containers, immersed in 6 liters of formalin) to prevent tissue decomposition. After three weeks of fixation, the hemispheres are transferred to and remain in air-tight MRI-compatible plastic containers fully immersed in 10% neutral buffered formalin. The brains are scanned in the same plastic containers to minimize tissue manipulation and prevent MRI artifacts caused by the presence of air bubbles in the ventricles and sulci due to exposure of tissue to air, which can in turn negatively impact the quality of the MRI acquisition, in particular for quantitative MRI sequences. The specimens that need to be scanned prior to the three-week timepoint are transferred to the same containers for scanning, and returned to the larger containers after the scan is completed. Tissue soaking in PBS is not performed as previous studies have reported inconsistent fluid penetration for whole brain specimens, creating artificial boundaries in the resulting images (Miller et al., 2011; Tendler et al., 2022).

The intact brain hemispheres are scanned using the 3T Siemens Prisma Fit human MRI scanner at the Douglas Cerebral Imaging Centre (CIC). The containers are stabilized in the 64-channel head/neck coil, with the hemispheres lying flat on the sagittal cut to minimize tissue motion and drifting during the scan and with the cerebellum towards in-bore to standardize the positioning and facilitate future application of automated image processing tools. The tissue is positioned in the center of the container (i.e. not touching the sides) to prevent potential artifacts. Figure 1 demonstrates the protocol for preparation of the tissue for scanning and positioning of the hemispheres in the 3T MRI scanner. Table 1 summarizes the 3T acquisition protocol used to acquire the postmortem images. All acquisitions were performed with one average.

**Table 1.** 3T MRI Acquisition Protocol.

| Sequence | TR (ms) | TE (ms) | FA (deg) | Matrix size | Resolution (mm<sup>3</sup>) | Bandwidth (Hz/Px) |
| --- | --- | --- | --- | --- | --- | --- |
| 3D T1w* | 2400 | 2.41 | 8 | 320×320×256 | 0.7×0.7×0.7 | 210 |
| 3D T2-SPACE | 2500 | 198 | 120 | 320×320×320 | 0.64×0.64×0.64 | 625 |
| 3D MGE-VFA | 38 | 4.27, 9.16, 14.05, 18.63, 22.82 | 7, 21, 35 | 160×160×256 | 1×1×1 | 210,210,210,24,250 |
| tfl_b1map | 197.80 | 2.41 | 8 | 96 × 96 × 220 | 2.3 × 2.3 × 3 | 490 |
| 3D meGSE (32 TEs) | 1000 | 9-288 | 180 | 128 × 123 × 230 | 1.8 × 1.8 × 1.8 | 1562 |
| GRE_fieldmap | 20 | 4.2, 9.90, 15.00 | 15 | 64 × 64 × 230 | 3.6 × 3.6 × 4.0 | 220,260,260 |
| DWI (64 directions) | 14000 | 101 | 90 | 70×128×128 | 2.0×2.0×2.0 | 2056 |

\*Magnetization Prepared RAPid Gradient Echo sequence, Inversion time = 1000 ms
TR: Repetition Time; TE: Echo Time; FA: Flip Angle; GRE: Gradient echo; DWI: Diffusion Weighted Imaging; meGSE: multi-echo Gradient and Spin Echo.

**Figure 1.**
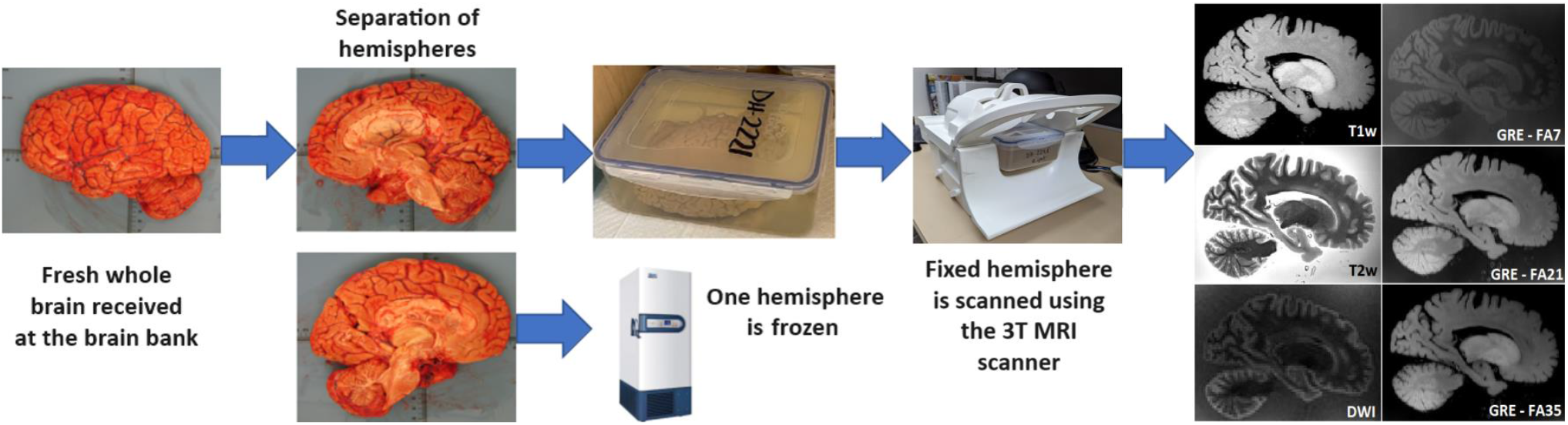
Flowchart demonstrating the preparation of the specimens for scanning. Hemispheres are separated by a sagittal cut in the middle of the brain, and are placed in an MRI compatible plastic container immersed in 10% formalin. The container is stabilized in the 64-channel head/neck coil, and scanned with the cerebellum in-bore.

A subset of the specimens is also scanned using an overnight 15-hour version of the same protocol, with higher resolutions and more averages, to provide additional neuroanatomical details. Table 2 summarizes the overnight 3T acquisition protocol.

**Table 2.** Overnight 3T MRI Acquisition Protocol.

| Sequence | TR (ms) | TE (ms) | FA (deg) | Matrix size | Resolution (mm<sup>3</sup>) | Averages | Bandwidth (Hz/Px) |
| --- | --- | --- | --- | --- | --- | --- | --- |
| 3D T1w* | 2400 | 2.59 | 8 | 384×384×224 | 0.6×0.6×0.6 | 8 | 210 |
| 3D T2-SPACE | 2500 | 160 | T2 var | 448×448×222 | 0.5×0.5×0.5 | 4 | 620 |
| 3D GRE (6 echoes) | 28 | 2.91, 5.41, ... 15.41 | 6**, 20 | 448×448×240 | 0.5×0.5×0.5 | 4 | 500 |
| tfl_b1map | 19780 | 2.41 | 8 | 96 × 96 × 220 | 2.3 × 2.3 × 3 | 1 | 490 |
| 3D T1-SPACE | 500 | 8.3 | T1var | 448×448×256 | 0.58×0.58×0.68 | 4 | 620 |
| 3D T1-IR*** | 2000 | 26 | 60 | 448×448×1854 | 0.4×0.4×0.4 | 2 | 266 |
| 3D meGSE (32 TEs) | 1000 | 9-288 | 180 | 128 × 123 × 230 | 1.8 × 1.8 × 1.8 | 4 | 1562 |
| GRE_fieldmap | 20 | 4.2, 9.90, 15.00 | 15 | 64 × 64 × 230 | 3.6 × 3.6 × 4.0 | 1 | 220, 260, 260 |
| DWI (64 directions)**** | 3000 | 90 | 90 | 138×221×221 | 1.6×1.6×1.6 | 1 | 2132 |

\*Magnetization Prepared Rapid Gradient Echo sequence, Inversion time = 1000 ms
\*\* One sequence with FA = 6 with magnetization transfer contrast (MTC) on.
\*\*\* The inversion time applied was 25 ms.
\*\*\*\* b-values from 1000 to 5000 s/mm<sup>2</sup>. Acceleration factor = 6
TR: Repetition Time; TE: Echo Time; FA: Flip Angle; GRE: Gradient echo; DWI: Diffusion Weighted Imaging; meGSE: multi-echo Gradient and Spin Echo.

Following the completion of the fixation process, smaller tissue blocks are extracted and scanned at higher resolution with an overnight 15-hour protocol using the 7T Bruker Biospec animal scanner at the Douglas CIC using comparable sequences. Table 2 summarizes the 7T acquisition protocol used to acquire the postmortem images. Consistent with the 3T protocol, the tissue blocks are scanned immersed in 10% formalin in MRI-compatible plastic containers. A 3D-printed holder is used to stabilize the container inside the 7T MRI coil to minimize tissue motion and drifting.

**Table S.2.**
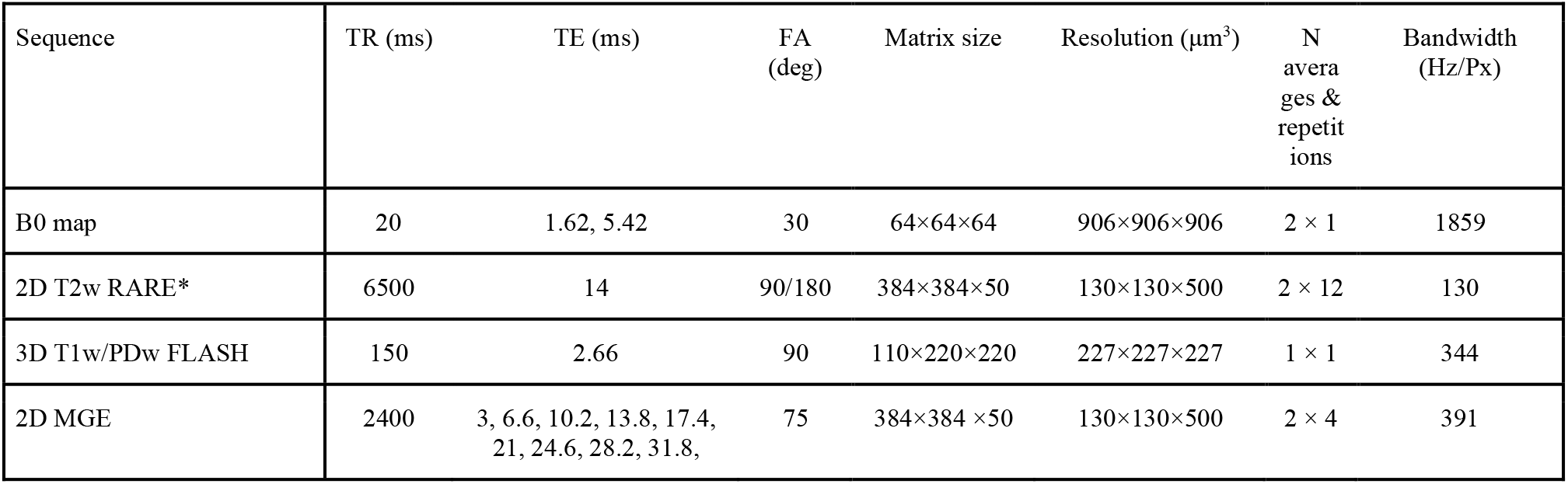
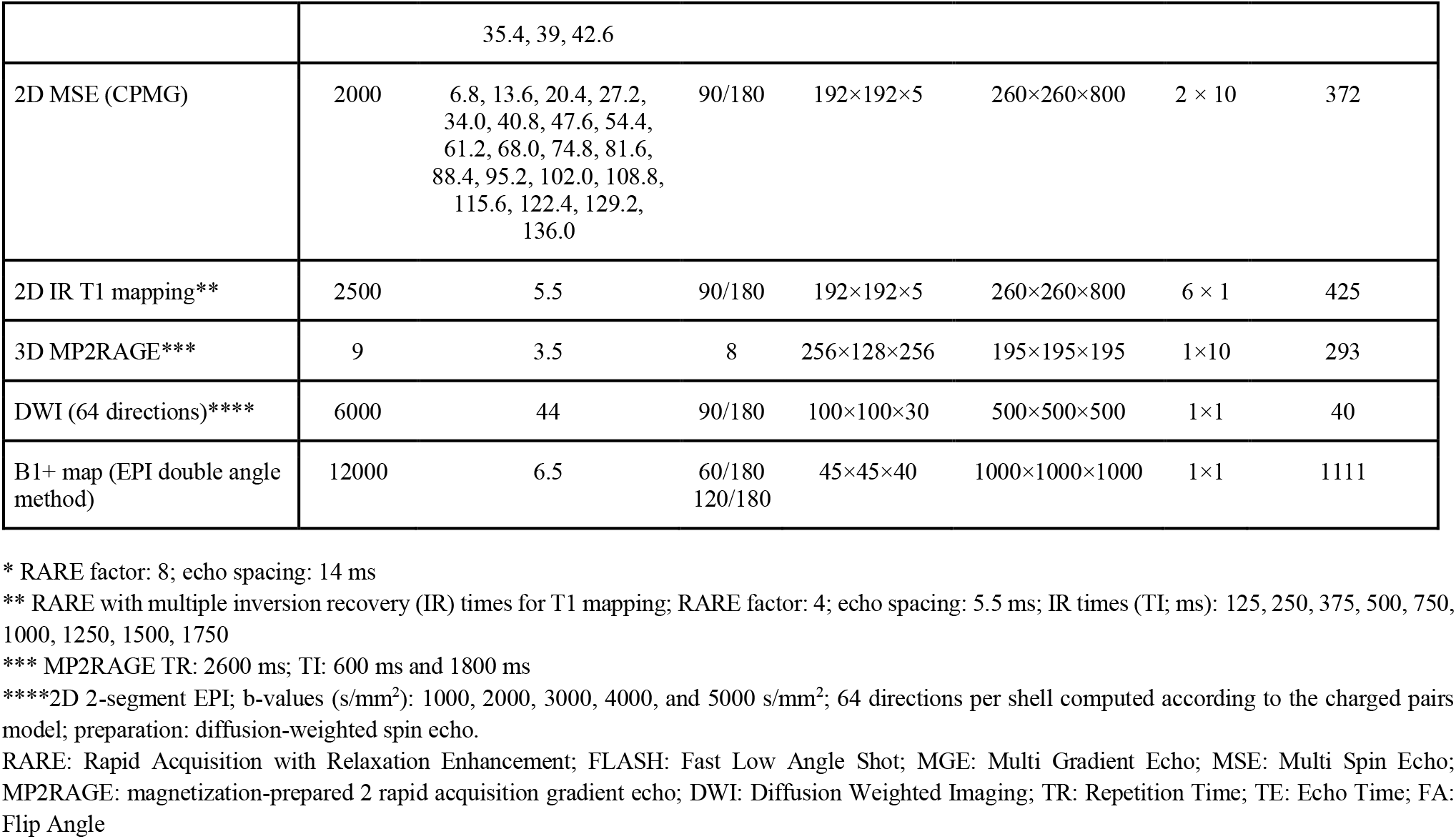
7T MRI Acquisition Protocol.

### Image Processing

The open access MINC toolkit V2 Version 1.9.18 (https://bic-mni.github.io/) was used to perform preprocessing, longitudinal co-registration of the T1w images, registration to stereotaxic space, and registration of the tissue block MRI to the hemisphere MRI (Dadar et al., 2018). BISON tissue classification tool was used to generate brain masks for quantitative and diffusion MRI processing (Dadar and Collins, 2021). To improve the processing of the diffusion images, two fast series of DWI with b = zero are acquired, using the DWI phase encoding (right-left) and the reverse (left-right). Mrtrix3 (Version 3.0.4) (Tournier et al., 2019) was used to denoise DWI volumes and to correct ringing artifacts. The mrtrix3 wrapper *dwifslpreproc* for FMRIB Software Library (FSL v6) (Andersson et al., 2003; Skare and Bammer, 2010) was then used to correct eddy-currents and susceptibility distortions, and the output was bias corrected using the ANTs N4 wrapper *dwibiascorrect* (Avants et al., 2009; Tustison et al., 2010). The tensor model (DTI) was then computed via Mrtrix3, and Fractional Anisotropy (FA) and Mean Diffusivity (MD) maps were extracted. We further performed probabilistic tractography on the preprocessed DWI volumes, seeding from the brainstem and terminating in the primary motor cortex using Mrtrix3’s *tckgen*. qMRLab was used to generate quantitative T1 and T2* maps (based on MGE-VFA acquisitions) as well as T2 and Myelin Water Fraction (MWF) maps (based on 3D meGSE acquisitions) from the quantitative MRI data (Duval et al., 2018).

## Results

### Sample Characteristics

The data presented in this section represents the characteristics of the postmortem images (N ∼ 200) acquired at the Douglas CIC between September 2022 and December 2023. To ensure the feasibility of the developed protocols across age ranges and disorders, postmortem scans were performed in brains with a range of disorders, including Alzheimer’s disease, Parkinson’s disease, Lewy body dementia, frontotemporal dementia, vascular dementia, mixed dementias, ALS, depressed suicide completers, as well as age-matched controls without any neurological disorders (mean age: 78.47 ±10.52, min age: 56, max age: 96, 44% females).

### T1 Weighted Images

As the process of fixation with formalin significantly impacts T1 relaxation times and consequently the gray-to-white matter contrast of the tissue in T1 weighted (T1w) MRIs (Pfefferbaum et al., 2004; Tendler et al., 2022), standard scans with Magnetization Prepared RApid Gradient Echo sequence (MPRAGE) were performed after a minimum of 4 months of fixation, to allow for the fixation process to complete for all brain regions, allowing for improved tissue contrast and signal to noise ratio (SNR). To verify the impact of fixation on different MRI contrasts, a subset of the specimens (N = 10) were also longitudinally scanned, from 0 to 120 days of fixation with 10% formalin. Figure 2 shows the impact of fixation on co-registered T1w MRIs of a DBCBB brain specimen that was scanned longitudinally from 0 days to four months of fixation. Note the changes in gray matter to white matter tissue contrast, noticeable at 3 days of fixation in the cerebellar and deep gray matter regions, expanding to the occipital lobe, and stabilizing in all brain regions after 90-120 days of fixation.

**Figure 2.**
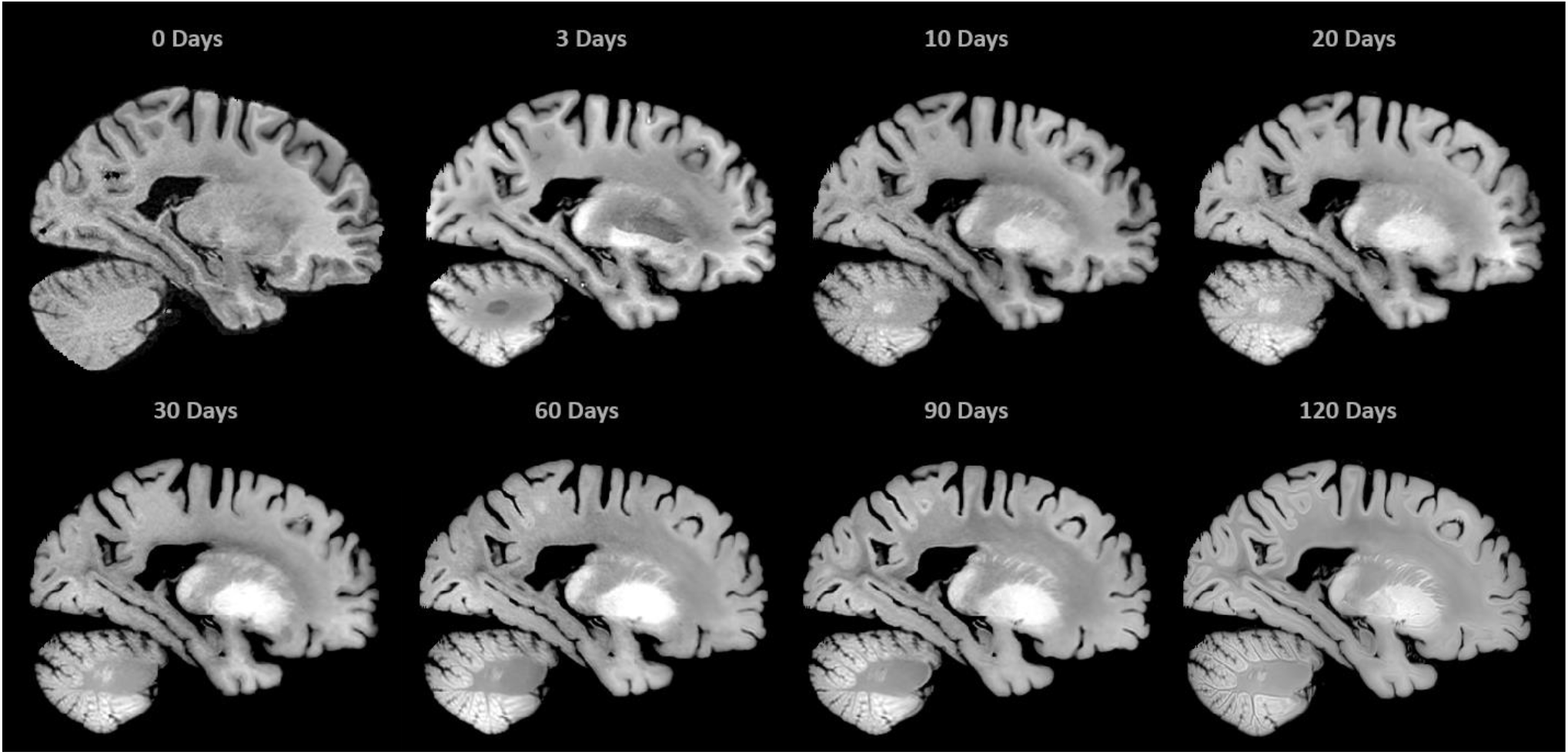
Change in T1w MRI contrast due to formalin fixation. Images show the same sagittal slices of one DBCBB brain scanned in 10% formalin from 0 days to four months of fixation. Note the contrast changes in cortical and deep gray matter regions, where after the completion of fixation, gray matter is hyperintense compared to the white matter. The 120-Day scan was acquired at higher resolution (the overnight protocol).

### T2 Weighted Images

While T2 relaxation times tend to decrease over long periods (years) of fixation (Dawe et al., 2014, 2009), the process of fixation does not significantly impact tissue contrast in T2 weighted (T2w) MRIs (Pfefferbaum et al., 2004; Roebroeck et al., 2019), and intensity normalization can standardize the overall intensity range of T2w images with different fixation times. Figure 3 shows intensity-normalized (linear normalization) sagittal slices of T2w images acquired from six different DBCBB specimens that were fixed from 0 days to 22 years, showing similar relative gray-to-white matter tissue contrasts to those observed in vivo T2w images. Note the relatively stable tissue contrasts for the long-term fixed tissues, allowing for use of a consistent protocol across different fixation times.

**Figure 3.**
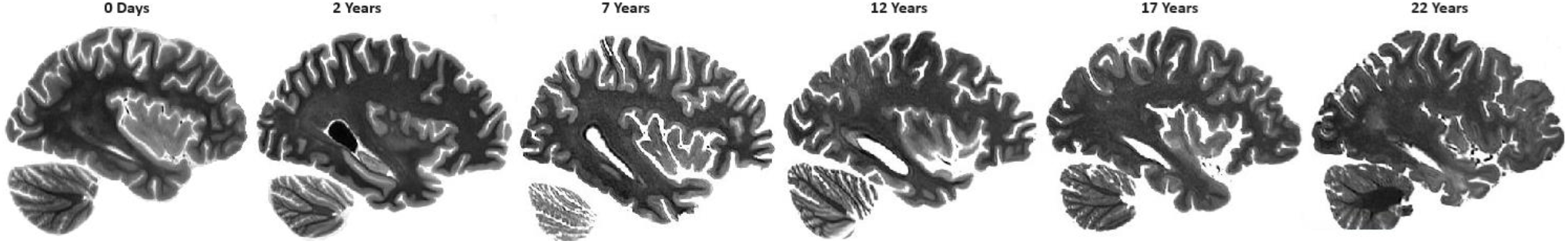
T2w images over long periods of fixation. Images show similar sagittal slices of six DBCBB brain specimens scanned in 10% formalin after fixation from 0 days to 22 years. Note that the gray to white matter contrast has remained relatively stable, and is consistent with in vivo T2w contrast.

In vivo, Fluid-attenuated inversion recovery (FLAIR) images are usually acquired for the detection of white matter hyperintensities (WMHs), since nulling of the cerebrospinal fluid (CSF) allows for easier differentiation of the WMHs from the CSF (Melazzini et al., 2021). We did not include FLAIR images in our postmortem protocol, as the fluid signal is no longer nulled for the formalin. As such, ex vivo FLAIR images do not provide additional information to the ex vivo T2w images, which can also be used to accurately detect and quantify WMHs both in vivo and ex vivo (McAleese et al., 2015; Dadar et al., 2017; Parent et al., 2023). See Figure 4 for examples of T2w and FLAIR images in three DBCBB specimens, where our standard T2w images are shown as the left hemispheres, and the corresponding FLAIR slices are flipped and shown as the right hemisphere to enable comparison. Also note the WMHs that are clearly visible on the T2w images (specimen shown on the right), with a high correspondence to the FLAIR-based WMHs.

**Figure 4.**
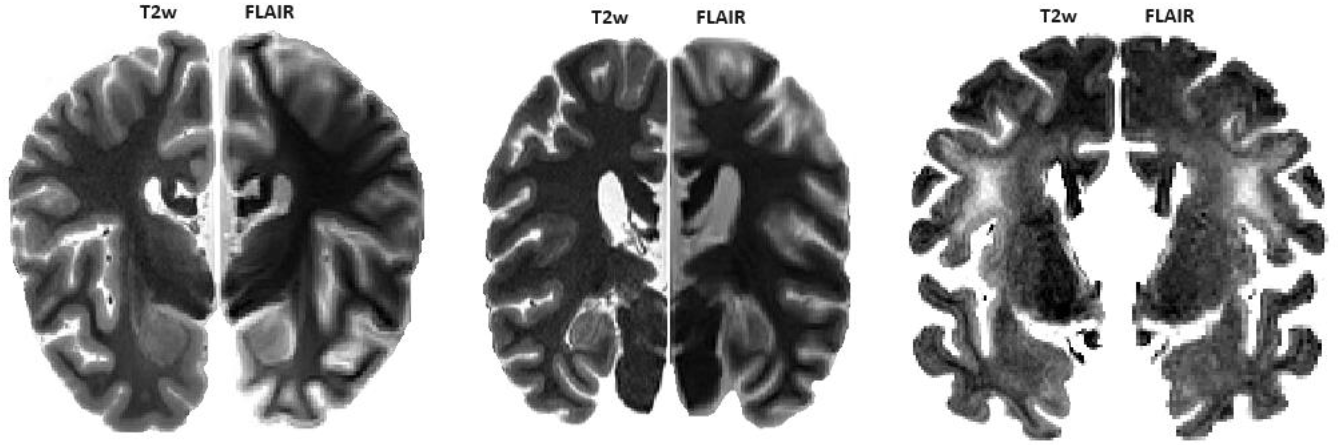
T2w and FLAIR images of three DBCBB specimens. Images show the same coronal slices of T2w (left) and FLAIR (right) images of the same specimens, showing similar hyperintense fluid signal for formalin. Note that the FLAIR images have lower resolution (1 mm^3^) than the T2w images (0.64 mm^3^).

### Diffusion Weighted Images

The process of fixation also significantly impacts the diffusion-weighted images (DWI), where T1, T2, and T2* relaxation times as well as diffusivity decrease in formalin-fixed tissues (Birkl et al., 2016; Dawe et al., 2009; Miller et al., 2011; Schmierer et al., 2008). Temperature is another factor that significantly impacts relaxation times and diffusivity in postmortem tissues, with mean diffusivity (MD) values reportedly decreasing by ∼30% when the brain temperature is decreased from body temperature (∼37 C) to room temperatures (∼20 C) (Berger et al., 2021). As such, the impact of temperature and fixation on diffusion metrics must be modeled and adjusted for, to enable comparison with in vivo metrics.

The overnight acquisition protocols include five b-values: 1000, 2000, 3000, 4000, and 5000. To find the optimal parameters for the DWI acquisition in the standard protocol, multiple DWI sequences were acquired for a subset of the specimens with a range of directions (30, 64, and 95) and b values (1000, 2000, 3000, 4000, and 5000). Based on these experiments, compared to including all five b-values, the b-value combination of 2000 and 5000 (64 directions) provided the optimal results in terms of SNR while keeping acquisition time feasible. Figure 5 shows an example of the calculated MD and FA maps for a DBCBB specimen with AD (fixation duration: 7 months), using all the acquired b values (right hemisphere) as well as using just b = 2000 and b = 5000 (corresponding slices, flipped and shown as left hemisphere). Performing probabilistic tractography on the preprocessed DWI volumes, we were also able to identify the corticospinal tract in both acquisition protocols as shown in the right panel (Figure 5).

**Figure 5.**
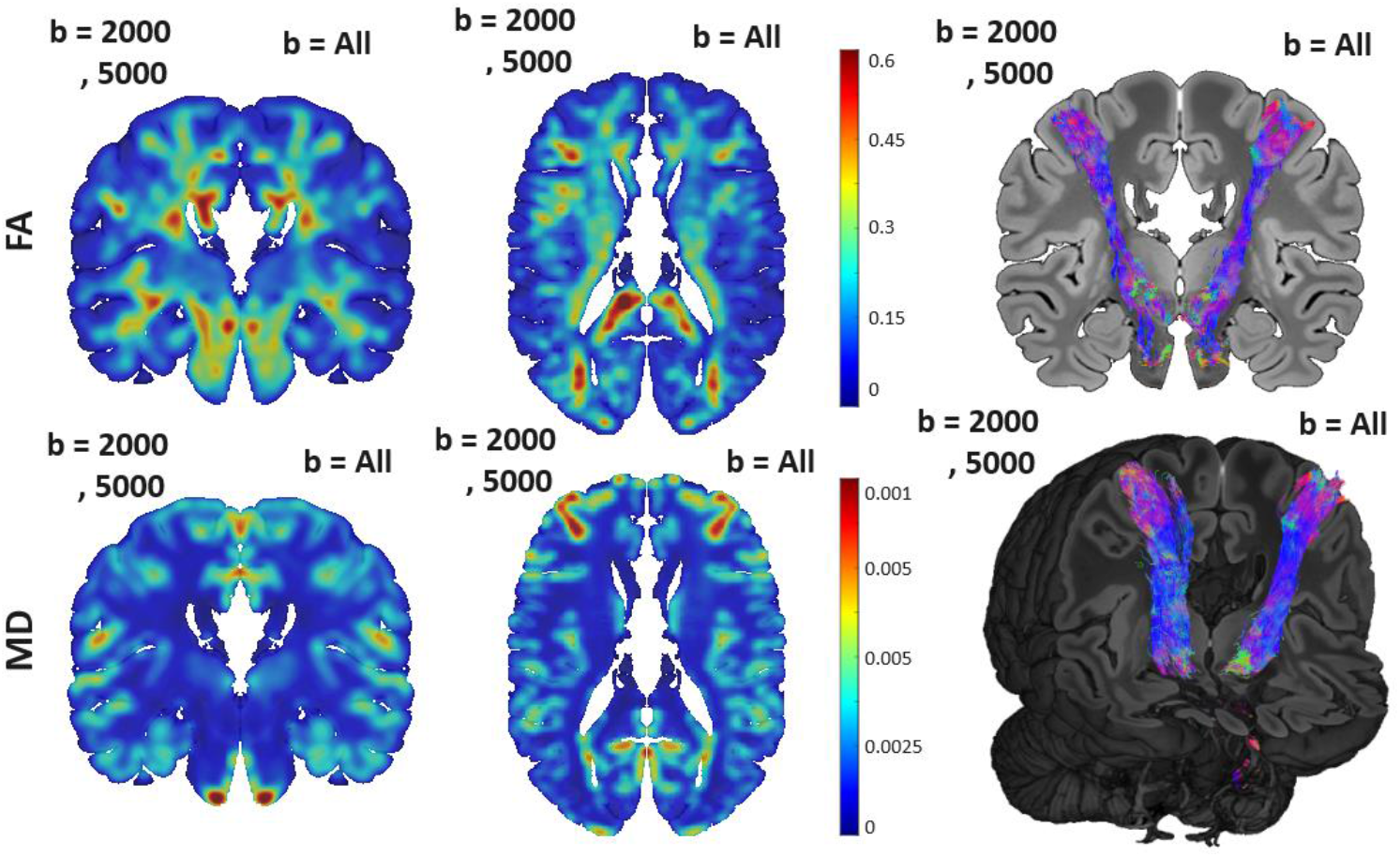
Left panel: Mean diffusivity (MD) and Fractional Anisotropy (FA) maps in white matter for one *DBCBB* specimen, estimated based on the ex vivo DWI data. Right panel: corticospinal tract identified through probabilistic tractography. Results shown on the right hemispheres are based on all 5 acquired b values, and results shown on the left hemispheres reflect the corresponding slices estimated using only b = 2000 and b = 5000.

### Quantitative Sequences

Three 3D gradient echo (GRE) sequences are acquired with different flip angles to calculate quantitative T1 maps (Fram et al., 1987; Marques et al., 2010). The variable flip angle (VFA) method with the addition of turbo-flash B1 map can be used to calculate T2 and T2* maps and perform quantitative susceptibility mapping (QSM). Furthermore, the currently used sequences that allow for T1 mapping free of bias fields to minimize inhomogeneity in transmit B1 (known as MP2RAGE) (Fram et al., 1987; Marques et al., 2010) did not provide sufficient contrast between gray and white matter on long term fixed brains. The 3D meGSE sequence acquires 32 TEs in 7:16 minutes and allows for fitting the multi-exponential model to generate MWF maps (Duval et al., 2018). As mentioned previously, since the process of fixation and to a lesser extent temperature impact T1, T2, and T2* relaxation times (Berger et al., 2021; Birkl et al., 2016; Dawe et al., 2009), the impact of temperature and fixation needs to be modeled and adjusted for, to enable comparison with in vivo quantitative metrics. Figure 6 shows the T1, T2, T2*, and MWF maps for one specimen. As expected and similar to previous reports (Dawe et al., 2009; Raman et al., 2017; Shatil et al., 2018), T1, T2, and T2* values are shorter compared to in vivo settings.

**Figure 6.**
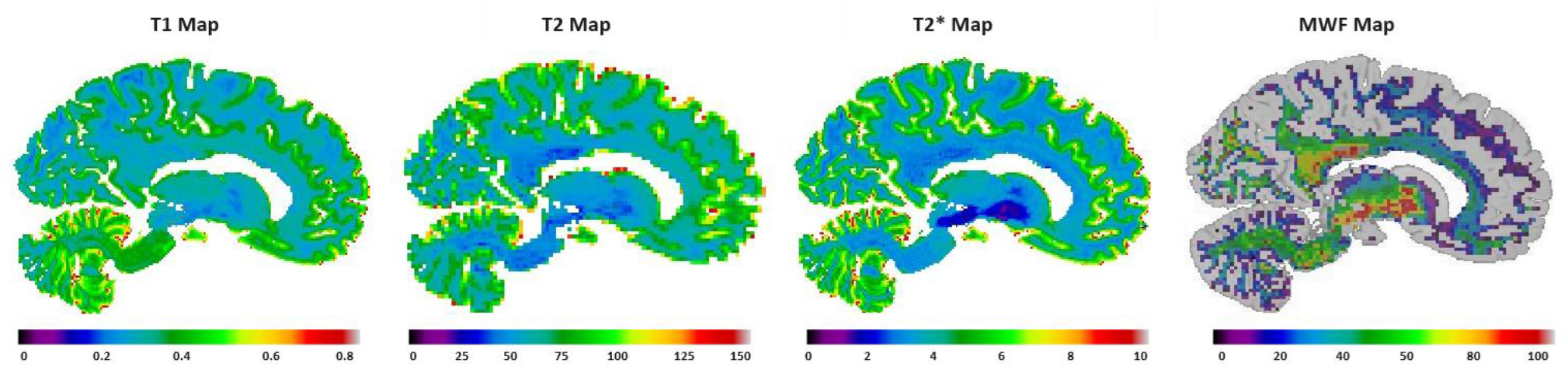
Quantitative T1, T2, T2*, and Myelin Water Fraction (MWF) maps for one DBCBB specimen were estimated based on the ex vivo GRE (T1 and T2* maps) and meGSE (T2 and MWF maps) data.

### Overnight 3T High-Resolution Scans

Figure 7 shows the co-registered standard resolution (coronal slice, shown as left hemisphere) and overnight high resolution (corresponding coronal slice, flipped and shown as right hemisphere) T1w (0.7 mm^3^ versus 0.5 mm^3^) and T2w (0.64 mm^3^ versus 0.5 mm^3^) images. Note that no preprocessing (i.e. denoising or intensity inhomogeneity correction) was performed on any of the scans, to allow for a fair comparison. The overnight scans achieve higher signal-to-noise ratio and significantly greater anatomical details as they are scanned overnight with hour-long acquisitions, allowing for better differentiation of subcortical gray matter structures and gray-to-white matter surfaces. This is particularly noticeable in the area of the external and extreme capsule where the claustrum can be precisely delineated in the high-resolution images, but not in the standard resolution ones. The nuclei of the subthalamic area, as well as the hippocampal folding and subsegments are also regions where the high-resolution images remarkably benefit the precision of the structures thanks to the decreased partial volume at the perimeters of these intricate brain nuclei and cortex regions.

**Figure 7.**
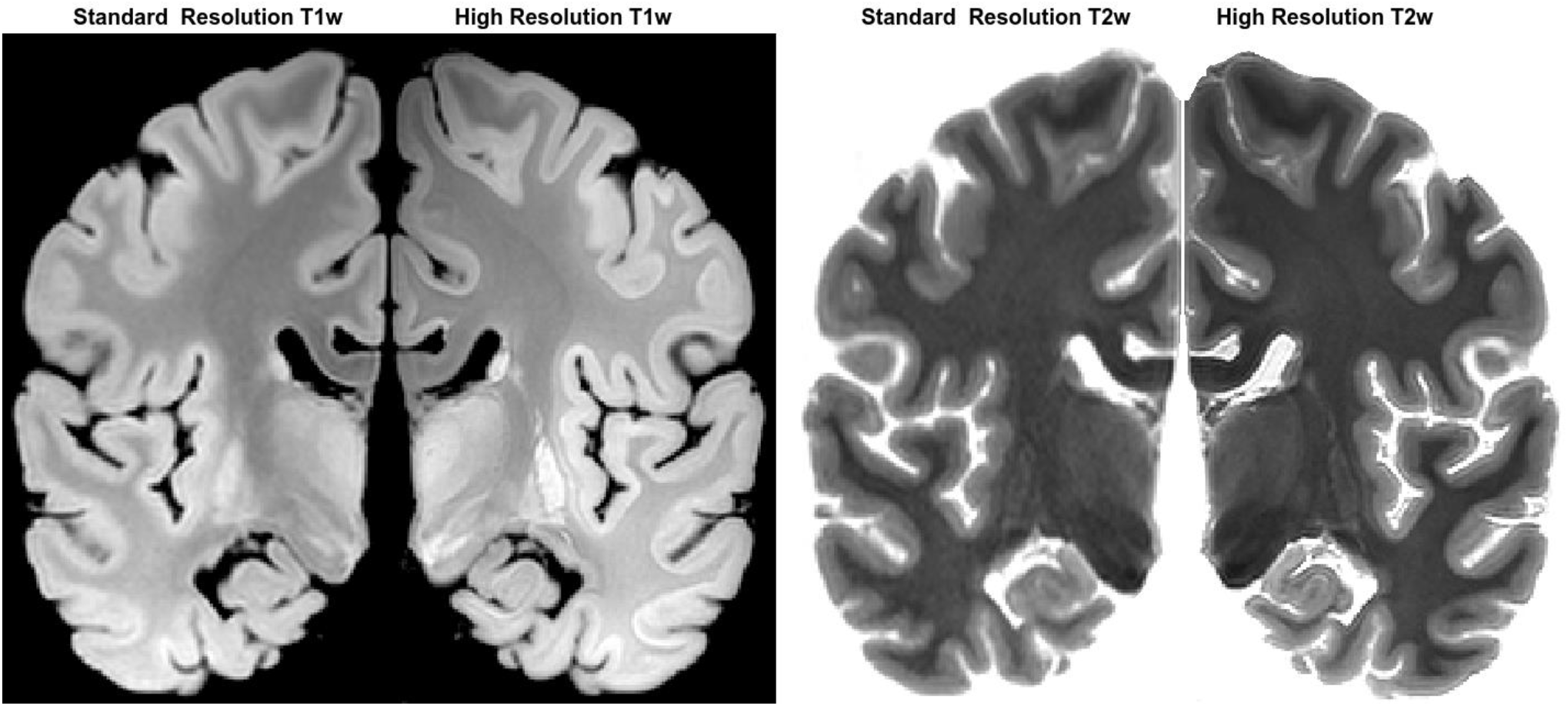
Corresponding standard resolution and high-resolution T1w and T2w images. Co-registered standard resolution (coronal slice, shown as left hemisphere) and overnight high resolution (corresponding coronal slice, flipped and shown as right hemisphere) T1w and T2w images.

### 7T Imaging of Tissue Blocks

Figure 8 shows co-registered 3T and 7T images of the same DBCBB specimen for a tissue block containing the left hippocampus. Note the high gray-to-white matter contrasted of both images as well as the added level of anatomical detail in the 7T images. Figure 9 shows the corresponding slices of T1w and T2w images of the same tissue block, as well as the derived T1 map, T2 map, and MWF results.

**Figure 8.**
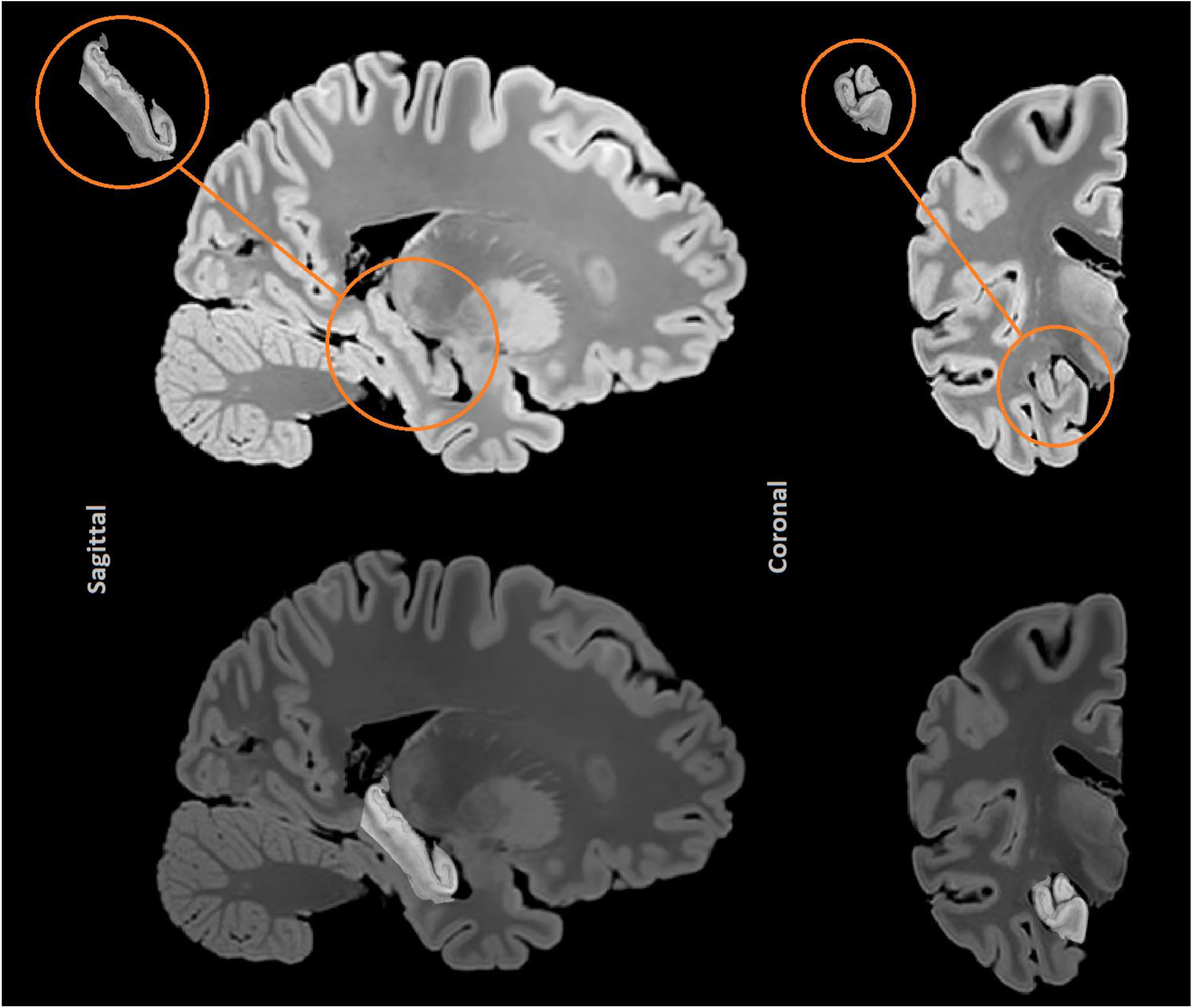
Co-registered 3T and 7T T1w images of a DBCBB specimen with histologically confirmed Alzheimer’s disease (fixation time: 12 years). Top row: corresponding sagittal (left) and coronal (right) slices of 3T and 7T T1w images. Bottom row: Sagittal and coronal 7T slices, overlaid on the 3T hemisphere images, demonstrating accurate registrations.

**Figure 9.**
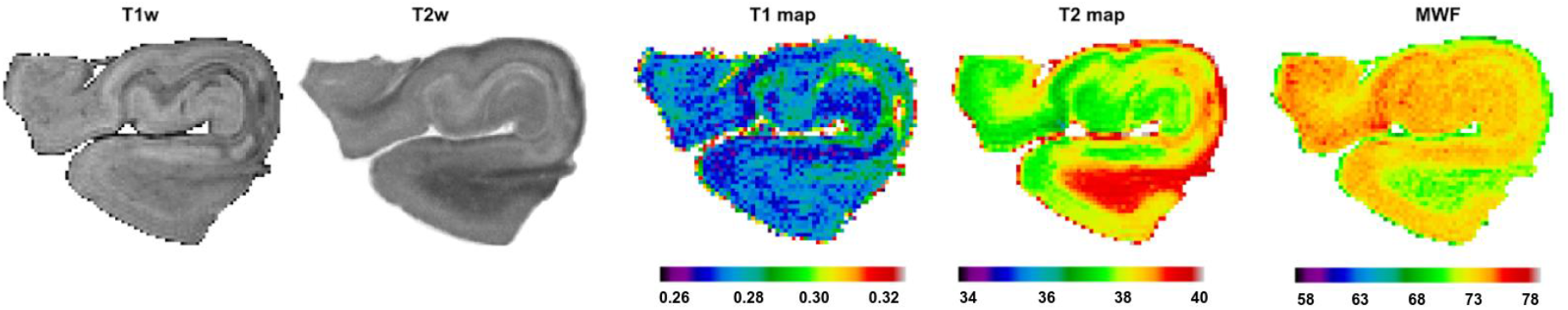
T1w, T2w, and quantitative T1, T2, and Myelin Water Fraction (MWF) maps for one DBCBB specimen (coronal slice, hippocampus) were estimated based on the ex vivo 7T T2 RARE (inversion recovery) and MSE data.

## Discussion

In this work, we present a post-mortem MRI protocol that has been developed for the DBCBB specimens, allowing for data acquisition from a large number of brains without jeopardizing tissue quality for follow-up histology and immunohistochemistry assessments. While performing large-scale harmonized and standardized in vivo imaging is a possibility for cohort studies of disorders such as neurodegenerative dementias (Belleville et al., 2019; Chertkow et al., 2019), this might not be the case for example for accidental deaths and/or suicide cases. Ex vivo MRI would therefore be an ideal method for obtaining anatomical and morphometric information of the intact brains in such cohorts before histology and immunohistochemistry assessments.

While post-mortem MRIs have been performed at smaller scales (Maranzano et al., 2020; McAleese et al., 2021; Miller et al., 2011; Pfefferbaum et al., 2004; Schumann et al., 2001; Tendler et al., 2022), large-scale high-resolution multi-modal post-mortem MRIs of intact brain specimens with a variety of neurodegenerative and mental health disorders has not been systematically performed. With our access to over 3000 post-mortem brain specimens at the DBCBB and the 3T and 7T imaging facilities at the Douglas CIC, we are able to perform ex vivo scans of hundreds of postmortem brains each year, building an invaluable repository of consistently acquired post-mortem MRIs of neurodegenerative and mental health disorders. To facilitate translation to in vivo and clinical settings, we aimed to acquire sequences that were similar or comparable to their in vivo counterparts.

The acquired high-resolution isotropic T1w and T2w 3T images with high contrast for gray and white matter allow for accurate volumetric and morphometric assessments of brain structures and quantification of abnormalities such as WMHs and tumors, whereas the DWI and quantitative sequences enable assessment of tissue microstructure and biophysical properties, once the impact of temperature and fixation are appropriately adjusted for. While additional processing of the postmortem tissue (e.g. soaking of the brains for improved diffusion, suspension in agar) might improve the quality of the postmortem MRIs in certain cases, they also introduce additional artifacts (e.g. intensity gradients and lines caused by inconsistent penetration of fluid, additional air bubbles trapped in the sulci and ventricles) (Tendler et al., 2022). Furthermore, they might impact the quality of the tissue for downstream histology and immunohistochemistry assessments and were generally not feasible for the large-scale of our ex vivo program. Instead, we chose to acquire a large number of consistently processed and scanned postmortem images with a large range of fixation durations (i.e. between 0 days to 22 years), allowing for future development of models that can be employed to adjust for the impact of fixation. (Miller et al., 2011)

Ex vivo DWI acquisition is particularly challenging due to a number of factors, including a decrease of mean diffusivity with a decrease in temperature (Berger et al., 2021), shortening of T2, and the consequent decrease in SNR (Miller et al., 2011). As such, ex vivo DWI acquisitions generally have poorer image quality and diffusion contrast compared to in vivo settings (Miller et al., 2011; Pfefferbaum et al., 2004). While modifications to in vivo protocols combined with longer acquisitions times (24 hours to 5 days) can lead to higher quality ex vivo DWI images (Miller et al., 2011), they were not feasible for our specific application. Instead, we acquired relatively lower resolution images (isotropic 2 mm^3^, 1.6 mm^3^, and 0.5 mm^3^ for the standard 3T, overnight 3T, and overnight 7T protocols, respectively). While the resulting MD and FA values were lower than their in vivo counterparts, we were able to retrieve the expected pattern in gray and white matter regions in MD, and the overall FA patterns within the white matter (Figure 5, left panel). Since the corpus callosum is cut by the sagittal split of the hemispheres, we used probabilistic tractography to identify the corticospinal tract as a quality control step (Figure 5, right panel).

The main challenge in ex vivo acquisition of the quantitative sequences pertained to the presence of small air bubbles in the ventricles and sulci that can lead to image artifacts, particularly in acquisitions with higher echo times and in the fresh tissue, where there was a greater likelihood for presence of air. In cases of prolonged fixation, air bubbles were naturally released in time, and we generally did not find air artifacts in the resulting images. In cases with shorter fixation periods (e.g. the longitudinally scanned specimens that had to be scanned from day 0), we slowly rotated the containers to release the air trapped in the sulci and ventricles. We further avoided the transfer of tissue across containers (except for the first three weeks of fixation, see the Methods section) to prevent the addition of further air bubbles.

In conclusion, high-resolution postmortem imaging can provide invaluable information on the intact brain before sectioning of the tissue for histology. The ex vivo images can complement information from in vivo imaging where in vivo imaging is possible, and provide novel information in the case of disorders where in vivo imaging might be challenging. A post-mortem imaging protocol that can be feasibly implemented for large-scale scanning of brain bank specimens is therefore of great value for researchers.

## Acknowledgements

Dadar reports receiving research funding from the Healthy Brains for Healthy Lives (HBHL), the Quebec BioImaging Network (QBIN), the Natural Sciences and Engineering Research Council of Canada (NSERC), and Brain Canada. Zeighami reports research funds from the HBHL and NSERC. Maranzano reports receiving funding from the QBIN and the NSERC.

